# SpaBatch: Batch Alignment of Spatial Transcriptomics Data using Graph Deep Learning

**DOI:** 10.1101/2025.03.24.645150

**Authors:** Jinyun Niu, Wenwen Min

## Abstract

Spatial transcriptomics (ST) has emerged as a transformative technology, enabling the simultaneous capture of gene expression and spatial context within tissues. However, batch effects arising from technical variability remain a significant barrier to integrating datasets generated from different experiments. To this end, we introduce a novel graph deep learning framework (SpaBatch) tailored for the batch alignment of multi-slice spatial transcriptomics data. SpaBatch is an innovative computational framework leveraging Variational Graph Autoencoders, self-supervised learning, and triplet learning with readout aggregation to enhance multi-slice spatial transcriptomics data integration. We validated our framework on multiple datasets encompassing various tissue types and experimental conditions. Through the adjustment of batch effects, SpaBatch facilitates the analysis of spatial transcriptome domains across multiple slices, showcasing its capability to reveal new biological insights in multi-slice spatial transcriptome studies.

## 1 Introduction

Spatial transcriptomics (ST) has emerged as a transformative technology, enabling the simultaneous capture of gene expression and spatial context within tissues. Multi-slice spatial transcriptomics (ST) data face challenges from batch effects caused by experimental condition variations, tissue sampling differences, biological heterogeneity, and developmental stage disparities across slices. Traditional single-slice clustering methods show limited performance in such data. To address these challenges, researchers have developed various batch correction tools. Harmony and Seurat v3, widely used integration tools in single-cell data analysis, have been adapted to spatial transcriptomics, leading to the development of tools like SEDR_Harmony and STAGATE_Harmony (Dong and Zhang, 2022; Korsunsky et al., 2019; Xu et al., 2024). However, directly applying these tools to ST data yields suboptimal results due to the more complex spatial architectures and cell-type variations in multi-slice datasets.

In this context, Zhou et al. (2023) developed STAligner, which calculates similarities between spots across slices using latent representations learned by STAGATE and aligns multi-slice data via the mutual nearest neighbors (MNN) algorithm while preserving spatial coherence. Although MNN-based methods excel in single-cell batch correction, they face challenges in multi-slice ST data: (1) Shared spatial structures across slices are not guaranteed, potentially leading to erroneous MNN pairings across distinct spatial domains; (2) Strict reliance on neighborhood relationships in latent space overlooks intra-domain anchor pairs not labeled as MNNs; and (3) Computational complexity escalates dramatically with increasing slice numbers, limiting scalability.

To further address batch effects, generative adversarial networks (GANs) have been introduced for multi-slice integration. Xu et al. (2023) proposed Splane, employing a multi-stage generation-adversarial strategy to eliminate inter-slice discrepancies. However, Splane exhibits subpar clustering performance with overly fragmented spatial functional regions. The SPIRAL framework from the University of Texas team incorporates gradient reversal layers (GRLs) to enhance adversarial training stability (Guo et al., 2023). While SPIRAL improves batch correction, its clustering results remain excessively scattered without biologically meaningful spatial aggregation. Additionally, Wang et al. (2023) developed STitch3D, which aligns 2D slices using iterative closest point (ICP) or PASTE algorithms, constructs global 3D neighbor graphs, and integrates slice/spot-specific effects with slice/gene-specific effects in a graph attention autoencoder model to mitigate cross-slice batch effects.

Despite innovative advancements in multi-slice analysis, existing tools still face limitations. However, some directly inherit single-cell methodologies without adequately addressing spatial transcriptomics-specific characteristics, while others struggle with computational efficiency for large-scale datasets. Current solutions thus require further optimization to address the complexity and batch effects inherent in multi-slice spatial transcriptomics data.

To this end, we propose a SpaBatch framework which is a novel computational framework designed for spatial transcriptomics (ST) data integration and analysis, particularly focusing on multi-slice datasets from diverse species, platforms, and tissue types. It combines Variational Graph Autoencoders (VGAE), self-supervised learning, and triplet learning with a readout aggregation strategy. Key innovations include masked data augmentation, k-nearest neighbor spatial graph construction, self-supervised deep embedded clustering (DEC), and triplet learning with readout aggregation, significantly improving clustering accuracy and cross-slice data integration.

We validate **SpaBatch** on seven spatial transcriptomics (ST) datasets spanning diverse tissues (human prefrontal cortex, mouse brain, embryonic heart, HER2+ breast cancer), species, and platforms (10x Visium, ST, Stereo-seq). Results demonstrate superior performance over state-of-the-art methods in spatial domain identification (average ARI = 0.613 on human DLPFC, 46% higher than STAligner) and batch effect correction (balanced iLISI/cLISI scores, outperforming methods like STG3Net and DeepST by 20–35%). SpaBatch uniquely resolves complex structures such as the hippocampal trisynaptic circuit in the mouse brain and tracks dynamic developmental changes in the human heart (4.5–6.5 PCW). Downstream analyses reveal spatially variable genes (e.g., *TMSB10* in cortical layers, *ERBB2* in breast cancer) and uncover tissue-specific biological processes via GO enrichment (e.g., immune response in tumor microenvironments). By integrating limited annotations in HER2+ breast cancer data, SpaBatch achieves semi-supervised clustering (median ARI = 0.367, 83% improvement over baselines), demonstrating robustness in pathological contexts. This work bridges critical gaps in multi-slice ST data integration, offering a unified framework for spatial omics studies and translational applications in developmental biology and cancer research.

## 2 Materials and methods

### 2.1 Datasets and data preprocessing

We applied SpaBatch to data from different species, platforms, and abnormal tissue slices for integrated analysis to validate the model’s performance.

The human dorsolateral prefrontal cortex (DLPFC) dataset measured by 10x Visium (Ji et al., 2020) came from three independent neurotypical adult donors, with each donor including four adjacent slices. The number of spots in each slice ranges from 3,498 to 4,789. Manual spot-level annotations from layer 1 to layer 6 and white matter (WM) provided by Maynard et al. (Maynard et al., 2021) were used as the ground truth.

The sagittal mouse brain data was analyzed using the 10x Visium platform, consisting of two groups: section 1 and section 2. Each section includes slices of the mouse anterior and posterior brain. Section 1 contains 2,695 spots in the mouse anterior brain and 3,355 spots in the mouse posterior brain. Section 2 contains 2,825 spots in the mouse anterior brain and 3,289 spots in the mouse posterior brain. Only the anterior brain slice of section 1 has manual annotations across all slices, which we used as the ground truth.

The coronal mouse whole brain dataset was provided by the ST platform, containing 35 coronal slices spanning the anterior-posterior (AP) axis (Ortiz et al., 2020). It includes manual annotations of 15 cluster types (Kleshchevnikov et al., 2022). These slices cover the entire brain region from the mouse olfactory bulb to the emergence of the cerebellum.

The early mouse embryo dataset described by Stereo-seq (Chen et al., 2022) included four slices from the E9.5 stage (E2S1, E2S2, E2S3, and E2S4), with 5,292, 4,356, 5,059, and 5,797 spots, respectively. Each slice has manual annotations.

The human embryonic heart dataset was profiled by the 10x Visium platform (Asp et al., 2019). It includes two sets of slices from human embryonic hearts at 4.5-5 PCW and 6.5 PCW. The 4.5-5 PCW period includes four slices, each with approximately 60 spots. The 6.5 PCW period includes nine slices, with spots ranging from 100 to 212.

The HER2-positive breast cancer dataset was profiled by the 10x Visium platform (Wu et al., 2021). The dataset includes tumor samples from 8 different patients, represented by groups A-H. Groups A-D each contain 6 slices, while groups E-H each contain 3 slices. Only the first slice in each group has manual annotations.

The input of SpaBatch consists of the gene expression matrices of multiple slices and their spatial locations. First, we concatenate the gene expression matrices along the spot dimension to construct a multi-slice gene expression matrix (hereafter denoted as the gene expression matrix). Genes expressed in fewer than 50 cells and with a total expression count less than 10 are removed. Subsequently, gene expressions are log-transformed and normalized based on library size using the SCANPY (Wolf et al., 2018) package. Finally, top-2,000 highly variable genes are selected, and Principal Component Analysis (PCA) is applied to reduce the dimensionality of the gene expression matrix while preserving as much data variability as possible, denoted as *X* ∈ *R*^*n×m*^.

### 2.2 Data augmentation with mask mechanism

Before training the model, we adopt a flexible masking mechanism to augment the pre-processed gene expression data. Specifically, we randomly sample a set of masked vertices *V*_*m*_ from all spots based on the masking rate *ρ*. If *V*_*i*_ (spot *i*) belongs to *V*_*m*_, then 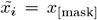, where *x*_[mask]_ denotes replacing the raw gene expression of spot *i* (*x*_*i*_) with a learnable vector; otherwise, 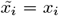. Therefore, the gene expression matrix *X* is re-defined as the masked gene expression matrix *X*_mask_ ∈ *R*^*n×m*^, defined as:

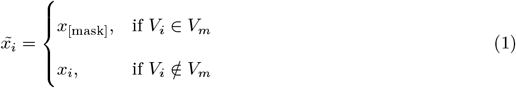

By adding a learnable vector to the masked spots, the model can dynamically adjust the representation of the masked spots during training, enabling them to learn features consistent with their surrounding neighbors. Through this mechanism, the model not only recovers the information of the masked spots but also enhances its generalization ability to the overall data.

### 2.3 Spatial graph construction

We use Euclidean distance to construct the spatial graph to represent the neighbor relationships between spots and select the *k*-nearest neighbors (Pedregosa et al., 2011). If spot *i* and spot *j* are neighbors, then *A*_*ij*_ = *A*_*ji*_ = 1. The adjacency matrix is symmetric and stored in the form of a sparse matrix. We can adjust *k* so that the number of neighbors for each spot ranges between 6 and 12, making it adaptable to different ST scenarios and platforms.

We calculate the neighbor relationships within each slice separately using the method described above, and then concatenate the adjacency matrices of each slice in a block diagonal form, denoted as *A* ∈ *R*^*n×n*^. It is to combine information from different slices to better capture the spatial relationships across slices.

### 2.4 Latent representation learning

The latent representation learning of gene expression is achieved through a variational graph autoencoder (VGAE), which consists of an encoder and a decoder (Figure 1). In the encoder, two fully connected layers are stacked to generate a low-dimensional representation 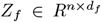 from the masked gene expression matrix *X*_[*mask*]_ ∈ *R*^*n×m*^. The Graph Convolutional Network (GCN) layer embeds the spatial graph *A* into *Z*_*f*_, capturing the spatial relationships among neighbors. The first GCN layer is used to learn a shared representation, following the approach of Kipf and Welling (Kipf and Welling, 2017), and is expressed as:

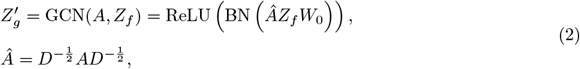

where *D* is the degree matrix, and BN stands for Batch Normalization. The second layer GCN formula is:

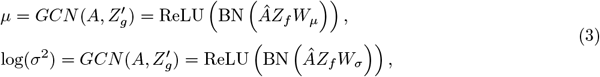

with two different parameters, *W*_*µ*_ and *W*_*σ*_, to independently model the mean and variance.

**Figure 1:**
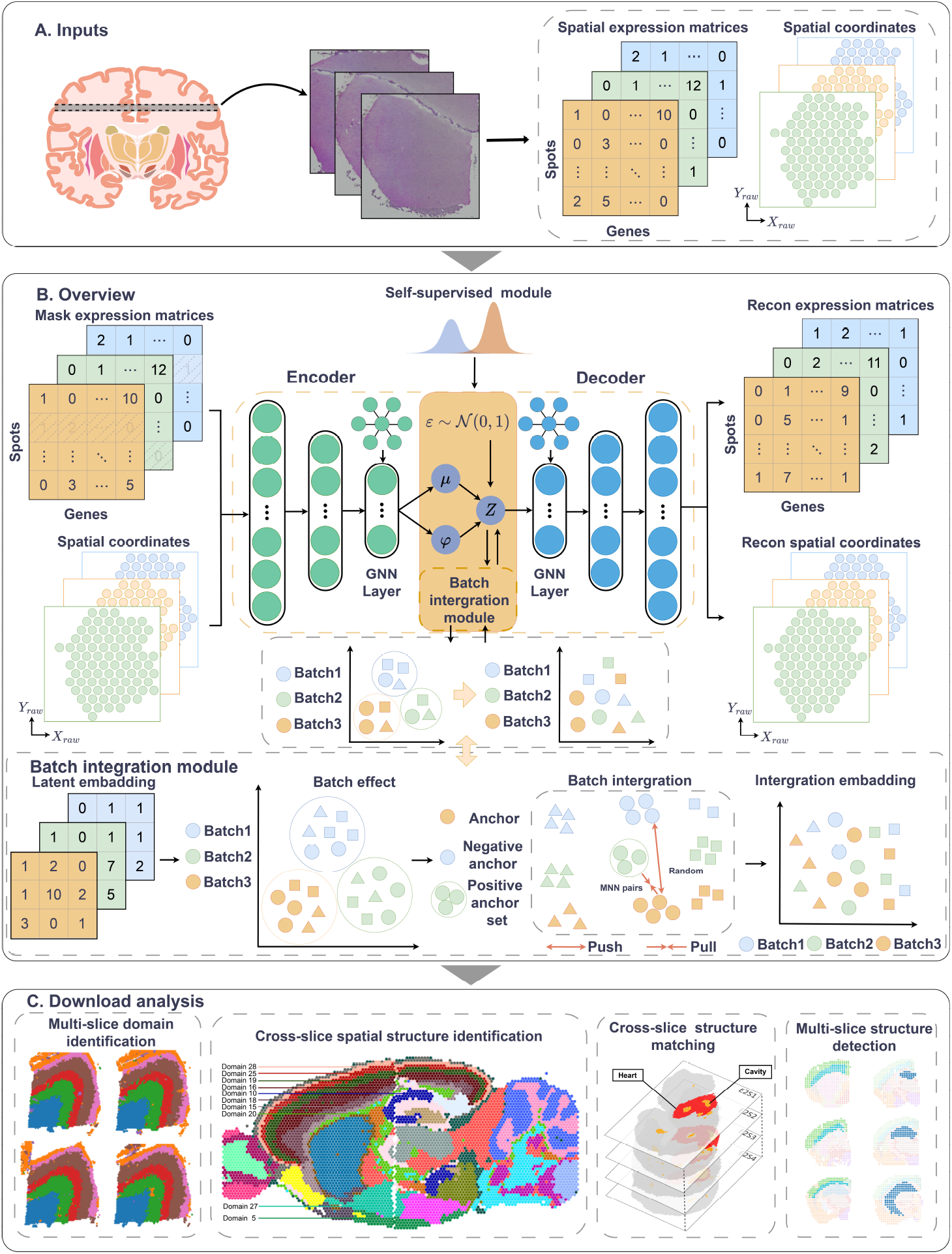
The architecture of SpaBatch. (A) Input data and data preprocessing. (B) Backbone network. (C) Downstream analysis.

Sampling directly from the distribution leads to non-differentiable gradients. To construct *Z*_*g*_, the reparameterization trick is employed:

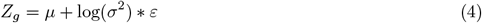

where *µ* and log(*σ*^2^) have been obtained from equation (3), *ε* is a randomly generated Gaussian noise. In SpaBatch, the final low-dimensional latent embedding *Z* = *Z*_*f*_ + *Z*_*g*_. After obtaining the low-dimensional latent embedding *Z*, an inner product decoder is used to reconstruct the adjacency matrix, denoted as *Ã*. To achieve this, the inner product decoder calculates the probability of an edge existing between each pair of nodes using the dot product of their respective latent vectors:

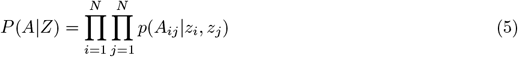

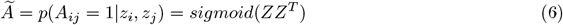

where *z*_*i*_ and *z*_*j*_ are the latent representations of spot *i* and spot *j*, respectively, and *sigmoid*(·) ensures that *Ã*_*ij*_ outputs a valid probability for the existence of an edge between spot *i* and spot *j*.

The objective of this reconstruction process is to approximate the raw adjacency matrix *A* by leveraging the learned latent representations. A loss function is constructed to minimize the binary cross-entropy loss between *A* and Ã:

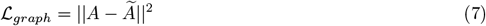

In addition to the reconstruction loss, the Kullback–Leibler (KL) divergence (Kingma and Welling, 2014) between the distribution of node representation vectors and a standard normal distribution is calculated. This term encourages the learned latent space to match a prior distribution. The KL divergence is given by:

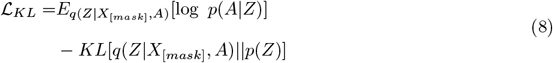

where 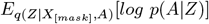 is the binary cross-entropy and *p*(*Z*) =Π _*i*_ *N* (0, *I*).

The decoder part employs a single layer of GCN to reconstruct the raw input gene expression matrix from the latent representation *Z*, denoted as 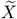. The GCN layer can capture local and global dependencies within the gene expression data, thereby assisting in the accurate reconstruction of the raw gene expression matrix. The reconstruction loss is constructed under a masked self-supervision framework by minimizing the difference between *X*_[*mask*]_ and 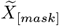. By employing the Scaled Cosine Embedding (SCE) loss as the objective function, it is formulated with a predefined scaling factor *γ* in the following manner:

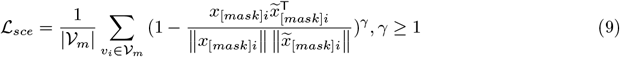

where *γ* is used to adjust the model’s sensitivity to larger errors. |*𝒱*_*m*_| is the number of spots in the masked set. *x*_[mask]*i*_ and 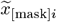 represent the feature vectors of the *i*-th spot in *X*[mask] and 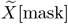, respectively.

#### 2.4.1 Self-supervision module

During the pretraining phase, SpaBatch learns the low-dimensional embeddings of gene expression in the latent space *Z* through a variational graph autoencoder (VGAE). Subsequently, deep embedded clustering (DEC) (Xie et al., 2016) is introduced to refine the local clustering details of *Z*. In the formal training phase, the model defines a clustering layer in the latent space, represented as 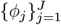, where *J* denotes the number of clusters. The self-supervised module first performs K-means clustering on *Z* and initializes the cluster centroids as the mean of samples in each cluster. These centroids are stored in the clustering layer and are further refined through iterative optimization to enhance clustering accuracy.

We use the Student’s *t*-distribution similarity (van der Maaten and Hinton, 2008) to quantify the relationship between spots and cluster centroids. Based on this similarity, it is transformed into the probability distribution *q*_*ij*_ that each spot belongs to a specific cluster, as defined by the following formula:

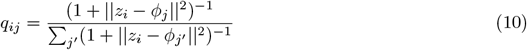

where *z*_*i*_ represents the embedding vector of spot *i* in the low-dimensional embedding *Z*, and *ϕ*_*j*_ corresponds to the *j*-th cluster centroid. The value *q*_*ij*_ is used to compute the soft assignment probability, describing the likelihood of spot *i* being assigned to cluster centroid *j*.

Additionally, the self-supervised module generates a target distribution by assigning higher weights to high-confidence samples (i.e., spots closer to the cluster centroids). This distribution is constructed by enhancing the peaks of the current soft assignment distribution, aiming to improve the model’s ability to distinguish between different clusters. The target distribution *p*_*ij*_ is defined as:

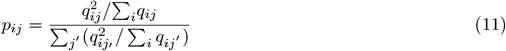

where *p*_*ij*_ represents the probability of assigning the *i*-th spot to the *j*-th cluster in the target distribution. The term ∑_*i*_ *q*_*ij*_ represents the total assignment probability for cluster *j*, normalized across all spots to ensure a valid probability distribution.

The self-supervised module minimizes the Kullback-Leibler (KL) divergence between the target distribution *p*_*ij*_ and the soft assignment probability *q*_*ij*_. The objective function is defined as:

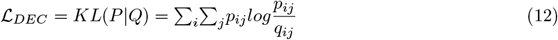

#### 2.4.2 Triplet learning based on readout aggregation strategy

To address the batch effect issue across multiple slices, we propose a triplet learning method based on the readout aggregation strategy (Long et al., 2023). Specifically, we first establish pairwise relationships between different slices and compute the spatial distances between spots in each slice and their paired spots in the other slice. This approach helps mitigate potential noise influences and enhances computational efficiency. When two spots from different slices are mutual nearest neighbors (MNN), we define them as anchor points. Then, we set a hyperparameter *α* to select the *α* nearest neighbors of the anchor points. These neighbors’ feature representations are fused into a positive anchor point using the readout aggregation function. In addition, the negative anchor point is randomly sampled to ensure feature differences between the anchor points. By aggregating the feature representations of the *α* neighbors, the readout aggregation function smooths the neighborhood information, making the generated positive anchor point more robust and reducing the impact of outliers in individual data points.

The triplet loss is employed to minimize the distance between anchor-positive pairs and maximize the distance between anchor-negative pairs in the latent space. The calculation is as follows:

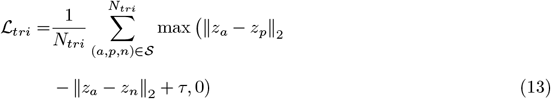

where *a, p*, and *n* represent the anchor point, the positive anchor point constructed using the readout aggregation strategy, and the negative anchor point, respectively. *N*_*tri*_ denotes the number of triplets in the set *S. τ* is the margin (with a default value of 1.0), ensuring that the distance difference between negative samples is sufficiently large.

### 2.5 Overall loss function

In the pretraining phase, we optimize only the three loss functions in VGAE: *ℒ*_*graph*_, ℒ_*KL*_, and *ℒ*_*sce*_, thereby obtaining the low-dimensional embeddings of gene expression in the latent space. In the training phase, we optimize the VGAE loss while updating the self-supervised module every 20 epochs and the triplet loss every 500 epochs, ultimately obtaining the final latent embeddings. The overall loss function is represented as:

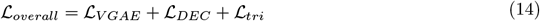

where *ℒ*_*V GAE*_ includes three loss functions in the backbone network: ℒ_*graph*_, *ℒ*_*KL*_, and *ℒ*_*sce*_.

### 2.6 Evaluation criteria

**ARI**.The Adjusted Rand Index (ARI) (Hubert and Arabie, 1985) is an external evaluation metric for measuring the similarity between two clustering results. It compares the relationship between the clustering outcomes and manual annotations, considering whether samples are correctly assigned to the same or different clusters. The range of ARI is [−1, 1], where values closer to 1 indicate better clustering performance, and values closer to 0 suggest that the clustering is similar to random assignment. The ARI is calculated as follows:

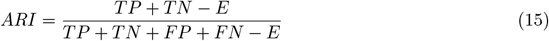

where *TP* is the true positive sample pair, *TN* is the true negative sample pair, *FP* is the false positive sample pair, *FN* is the false negative sample pair. *E* is the expected similarity, it is calculated as follows:

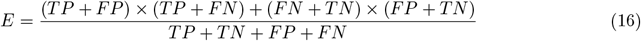

**ACC**.Normalized Mutual Information (NMI) measures the amount of shared information between two clustering results. The value of NMI ranges from [0, 1], with larger values indicating better clustering. Adjusted Mutual Information (AMI), on the other hand, is an adjusted version of mutual information that removes the influence of randomness, and its value ranges from [−1, 1], with larger values indicating better clustering (Yuan et al., 2024). We combine NMI and AMI to assess the consistency of clustering results, denoted as Average Clustering Consistency (ACC). The specific formula for ACC is as follows:

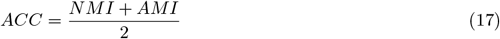

ACC can comprehensively consider the similarity and information sharing between clustering results, thus providing a thorough evaluation of the clustering performance.

**V-measure**.V-measure is a clustering evaluation metric based on information theory (Hu et al., 2024). It consists of two components: Homogeneity (HOM) and Completeness (COM). V-measure calculates the harmonic mean of these two metrics to comprehensively assess the consistency and completeness between the clustering results and the manual annotations, thereby evaluating the quality of the clustering. The value of V-measure ranges from [0, 1], with a higher value indicating better clustering results. Its calculation formula is as follows:

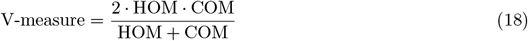

**LISI**.Integration Local Inverse Simpson’s Index (iLISI) and Clustering Local Inverse Simpson’s Index (cLISI) are two clustering performance evaluation metrics based on the Inverse Simpson’s Index, commonly used to measure the success of batch effect correction in multi-batch datasets (Tran et al., 2020). iLISI measures the effective number of different batches within a local neighborhood of a cell. A higher iLISI value indicates better batch effect correction, meaning that even after batch correction, cells from different batches but with similar biological characteristics can still cluster together in the latent space. cLISI measures the effective number of different cell types within a local neighborhood of a cell. A lower cLISI value indicates better multi-slice integration, reflecting that cell types from different batches have been successfully integrated and batch effects have been effectively corrected.

### 2.7 Implementation Details

In this study, all experiments were conducted on a single NVIDIA RTX 4090Ti GPU. The mask rate was set to 0.2. We adjusted the parameter *k* used to construct the spatial graph, setting the number of neighbors between 6 and 12, making it adaptable to different ST scenarios and platforms. According to our tests, when *k* is set to 8, SpaBatch achieves the best performance across most datasets. The fully connected layers of the encoder had dimensions of 64 and 16, while the graph convolution layers were set to 64 and 16. The number of clustering centers in the self-supervised module was set to 20. The learning rate and weight decay were set to 5e-4 and 1e-4, respectively, and optimization was performed using Adam.

### 2.8 Baseline methods

We compared SpaBatch with state-of-the-art methods. Here are the descriptions of the methods and the parameter settings:

- **STAligner** uses STAGATE as the backbone network to learn low-dimensional embeddings of gene expression and constructs MNN pairs for training in the latent space. The data preprocessing was performed using the default parameters, with top-5000 highly variable genes selected as input. The “rad_cutoff” parameter is adjusted across different datasets to ensure the number of neighbors is between 6 and 12, achieving optimal performance.
- **STG3Net** uses a masked graph convolutional autoencoder as the backbone module, combined with generative adversarial learning and a global neighbor selection strategy to construct triplets for robust multi-slice spatial domain identification and batch correction. We used the default parameter settings.
- **SEDR** is a graph neural network-based spatial embedding method. SEDR applies Harmony (Korsunsky et al., 2019) to perform batch correction on the learned low-dimensional embeddings. We followed the parameter settings provided in the authors’ code, specifically setting “using_dec” to False and the number of “epochs” to 200.
- **DeepST** extracts features from H&E images based on spatial location information and constructs a neighbor graph from the image features to enhance gene expression. Since DeepST requires H&E images as input, we applied it only to the DLPFC and sagittal mouse brain datasets. We set the default parameters according to the demonstration code in the online tutorial.
- **SpaGIC** learns meaningful point latent embeddings by maximizing the mutual information between edges and local neighborhoods of the graph structure and minimizing the embedding distance between spatially adjacent points. It integrates graph convolutional networks and self-supervised contrastive learning techniques. SpaGIC uses Harmony to batch-correct the learned low-dimensional embeddings. We set “mse_weight”, “graph_weight”, and “nce_weight” to 60, 0.01, and 0.01, respectively. The number of “epochs” was set to 500, and “n_neighbor” was set to 5, all consistent with the default parameters.
- **STitch3D** leverages the ICP or PASTE algorithm to optimize the alignment between multiple slices. It incorporates both slice-spot and slice-gene factors, and reconstructs gene transcription expression by utilizing cell composition components informed by scRNA-seq data.

## 3 Results

### 3.1 SpaBatch effectively corrects batch effects and precisely identifies spatial domains in the DLPFC dataset

To quantitatively evaluate SpaBatch in spatial domain identification and batch effect correction for multi-slice joint analysis, we first applied it to the human dorsolateral prefrontal cortex (DLPFC) dataset measured by 10x Genomics Visium (Ji et al., 2020). The data consist of 12 slices from three independent neurotypical adult donors, with each slice containing four adjacent slices (Figure 2A). Maynard et al. (Maynard et al., 2021) provided manual spot-level annotations from layer 1 to layer 6 and the white matter (WM), which were used as ground truth in the evaluation. We compared SpaBatch with six state-of-the-art methods, evaluating the spatial domain identification performance of each method using the Adjusted Rand Index (ARI) (Hubert and Arabie, 1985), Average Clustering Consistency(ACC) and V-Measure. Additionally, we assessed batch effect correction and multi-slice integration results using the Integration Local Inverse Simpson’s Index (iLISI) and Clustering Local Inverse Simpson’s Index (cLISI) (Tran et al., 2020).

**Figure 2:**
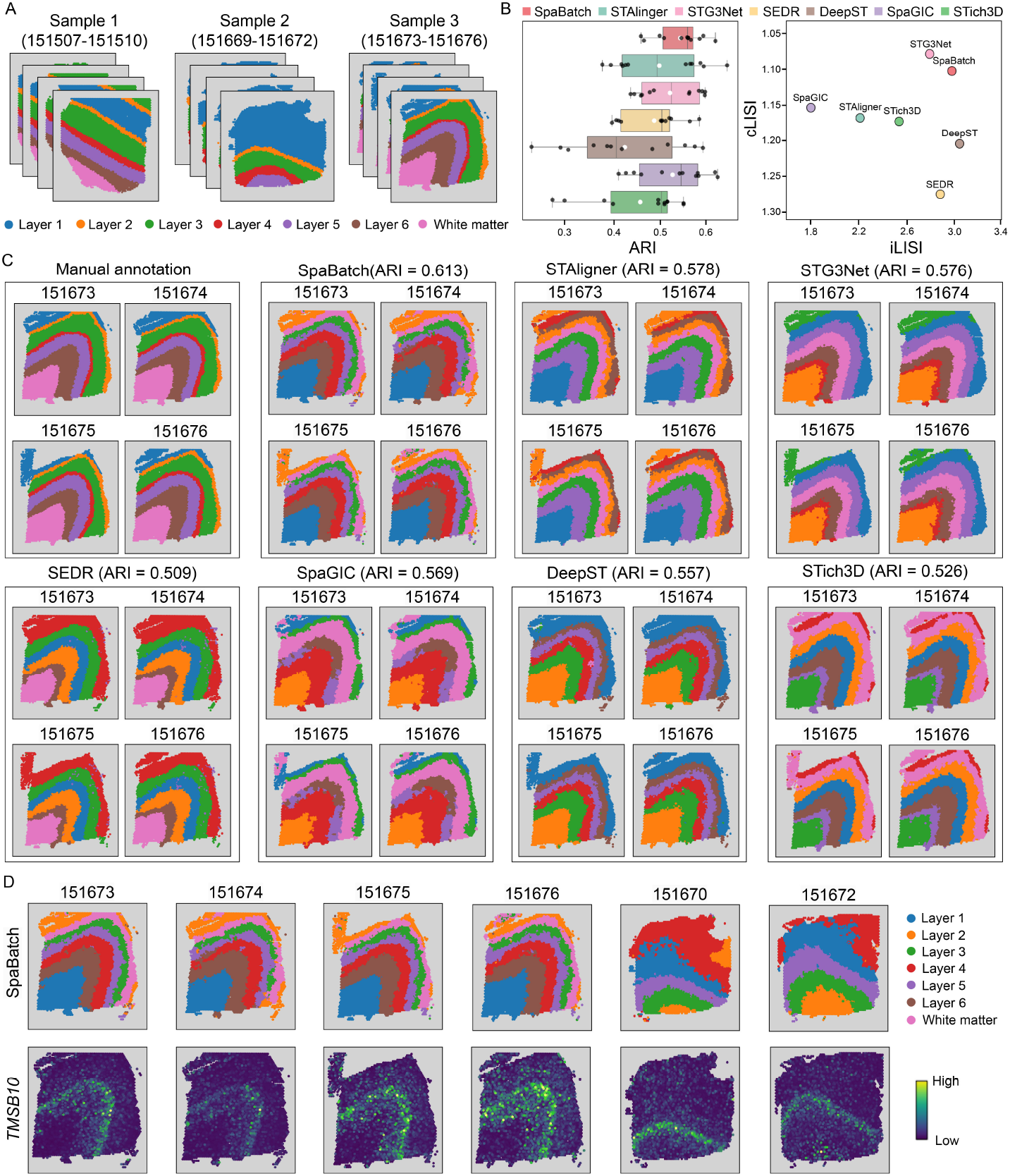
SpaBatch and other methods for multi-slice joint analysis on the DLPFC dataset. A. Three samples of the DLPFC dataset and their manual annotations. B. Boxplots of ARI values calculated by SpaBatch and other methods across 12 slices in three samples of the DLPFC dataset. In the boxplots, the central line and the solid white dot represent the median and the mean, respectively. The swarm plot illustrates the accuracy distribution across all slices (left). The iLISI and cLISI scores calculated for SpaBatch and other methods on three samples of the DLPFC dataset are shown (right). The x-axis represents batch mixing scores, and the y-axis represents spatial domain mixing scores. Points closer to the top-right corner indicate better performance. C. Integration results of four slices from sample 3 of the DLPFC dataset by SpaBatch, with identification of spatial domains. D. Spatial domains identified by SpaBatch across six slices (top) and the spatial expression of the layer 5 marker gene *TMSB10* (bottom).

We divided the samples from different donors into three groups, with each sample containing four adjacent slices (Figure 2A). We first evaluated the performance on sample 3 and observed that the spatial domains identified by SpaBatch, STAligner, and STG3Net were well-mixed within the same cortical layer, while different cortical layers were ordered according to the spatial structure of layer 1 to layer 6 and the white matter layer (WM). Compared to the latter two methods, SpaBatch produced more precise boundaries and shapes. Other methods exhibited inaccuracies in spatial domain identification within layers 1-4. SpaBatch achieved the highest clustering accuracy in terms of ARI, with an average value of 0.613, surpassing STAligner (ARI = 0.578) and STG3Net (ARI = 0.576), and significantly outperforming the other methods (Figure 2C). To further validate, we conducted the same experiment on sample 1 and sample 2. The results show that SpaBatch continues to achieve the spatial domain identification results closest to manual annotation (Supplementary Fig. S1). We conducted experiments on samples 1-3 and calculated the ARI values for the 12 slices. The results indicate that SpaBatch performed the best, achieving the highest median and mean values, further emphasizing its advantage in spatial domain identification on the DLPFC dataset compared to other methods (Figure 2B (left) and Supplementary Fig. S2A). In addition, we further tested the robustness of SpaBatch by comparing the clustering accuracy across different random seeds and found that SpaBatch is not sensitive to variations in random seeds (Supplementary Fig. S2B).

The Uniform Manifold Approximation and Projection (UMAP) (Becht et al., 2019) visualizations created using Scanpy (Wolf et al., 2018) clearly indicated that the layer structures of these methods are well-organized and highly consistent with the manual annotations. In particular, SpaBatch demonstrated an exceptional ability to accurately capture the spatial domain structure, with the UMAP visualization closely resembling the manual annotation of layer distribution. Additionally, SpaBatch excelled in achieving smooth and uniform mixing across different batches, as seen in the batch integration across slices. This stood in sharp contrast to other methods, such as SpaGIC and STAligner, which exhibited uneven mixing in certain local regions (Figure 3). The values of iLISI and cLISI also indirectly demonstrated the ability of SpaBatch to achieve effective slice mixing and batch effect correction. While the higher cLISI score of STG3Net highlighted its strength in batch effect correction, it came at the expense of reduced iLISI, indicating potential loss of biological signals. On the other hand, the higher iLISI score of DeepST reflected better preservation of biological signals but revealed poor performance in cLISI, suggesting insufficient batch effect correction. SpaBatch demonstrated a balanced performance, excelling in cLISI while maintaining a high iLISI score. This balance showcased the ability of SpaBatch to perform batch effect correction effectively while preserving critical biological signals (Figure 2B (right)). SpaBatch not only achieved higher accuracy in spatial domain identification but also ensured that the integration across batches maintained a high degree of consistency and coherence. This advantage continued to hold across other samples (Supplementary Figs. S3 and S4).

**Figure 3:**
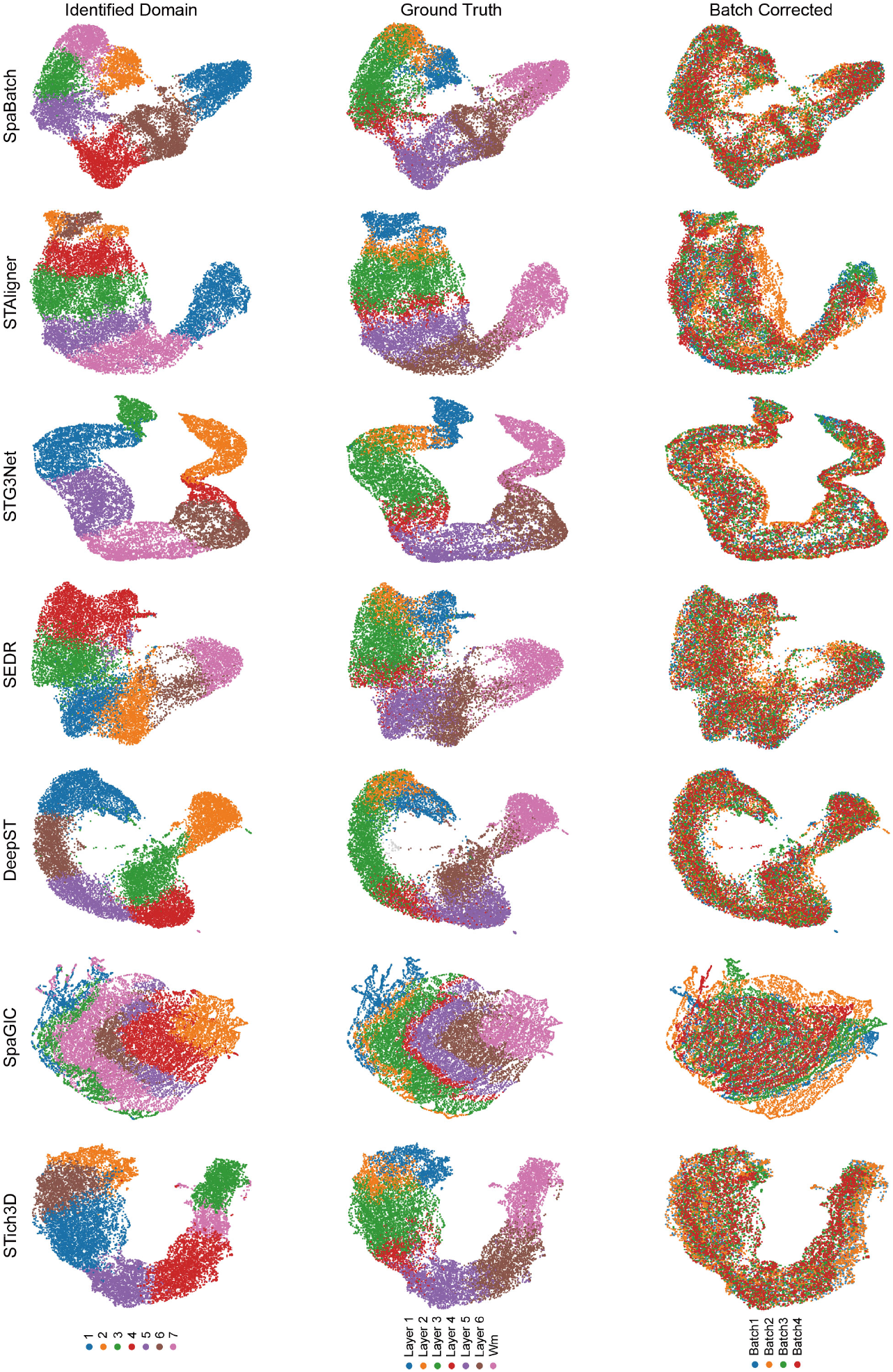
UMAP visualization of donor3 embeddings colored by identified domains (left), ground truth (middle), and batch corrected (right).

We further defined layer-marker genes through differential analysis and previous reports (Maynard et al., 2021), such as *AQP4* (layer 1), *CARTPT* (layer 2), *ENC1* (layer 3), *PCP4* (layer 4), *TMSB10* (layer 4 and layer 5), and *MBP* (WM), which exhibited significant expression differences across different cortical layers (Supplementary Fig. S5). By visualizing the layer-marker gene *TMSB10* in layer 5, we demonstrated that SpaBatch can effectively identify layer-marker genes and depict the shared organizational structures between different samples (Figure 2D).

### 3.2 SpaBatch comprehensively depicts the mouse brain from both macro and micro perspectives

Next, we evaluated the performance of SpaBatch on more complex tissue sections. We conducted experiments on sections 1 and 2 of sagittal mouse brain data (Figure 4A and Supplementary Fig. S6A, S7A) generated using the 10X Visium protocol (Ji et al., 2020). Both sections 1 and 2 contain paired samples of the anterior and posterior brain. We integrated the anterior and posterior regions of sections 1 and 2 separately for the experiments. Among the two datasets, only the anterior region of section 1 has manual annotations, which we used as ground truth (Supplementary Fig. S6B). The data provided an overall anatomical structure for understanding the mouse brain’s organization from a global perspective.

**Figure 4:**
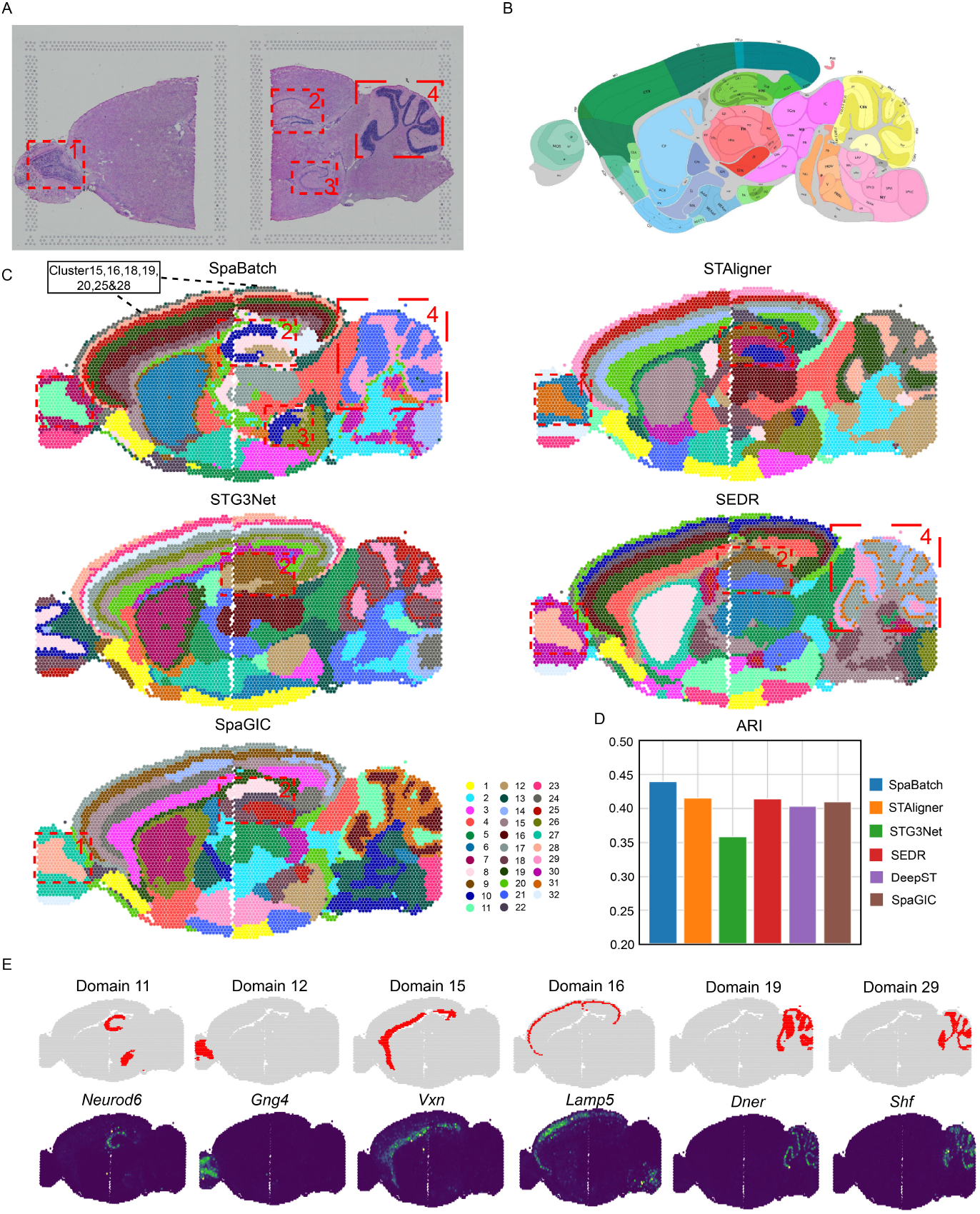
SpaBatch comprehensively depicts the sagittal mouse brain from the macro perspective. A. H&E images of sagittal anterior and posterior sections of Section 1, and the corresponding specific spatial subdomains. B. Manual annotation of the sagittal anterior mouse brain (Section 1). C. The spatial domain identification results of sagittal mouse brain Section 1 integrated by SpaBatch and other methods. We used red boxes and numbers to highlight specific spatial subdomains. D. The bar plot comparing the ARI values obtained from SpaBatch and other methods with the manual annotation. E. SpaBatch identifies six distinct fine spatial subdomains (top) and the spatial expression of marker genes associated with these subdomains (bottom).

We first tested the integration capabilities of SpaBatch, STAligner, STG3Net, SEDR, and SpaGIC using Section 1. For comparison, the number of clusters for all algorithms was set to 32. The mouse brain atlas provided by Allen Brain Atlas (Sunkin et al., 2012) was used as a reference (Figure 4B). We found that SpaBatch, STAligner, STG3Net, and SEDR were all able to identify shared clusters between adjacent regions of consecutive slices in section 1 and section 2, primarily including the cortex, hippocampus, thalamus, and hypothalamus. For more delicate tissue structures, SpaBatch has a stronger detection capability. The main olfactory bulb in the anterior and the cerebellar cortex in the posterior (red box 1 and 4 of Figure 4C)) detected by SpaBatch show high consistency with the Allen Brain Atlas mouse cortex reference atlas. Only SEDR identified both spatial domains simultaneously, but its accuracy in the cerebellar cortex region was still inferior to SpaBatch. Moreover, we further examined the dorsal and ventral regions of the hippocampus located in the posterior brain (red box 2 and 3 of Figure 4C). The dorsal hippocampal region consists of CA1, CA2, CA3, and the dentate gyrus (DG), which together form the classic trisynaptic circuit. This circuit is crucial for memory formation and retrieval (Rotenberg et al., 1996). Among all methods, only SpaBatch accurately identified the ventral region of the hippocampus and fully recognized the ammonic horns (CA), composed of CA1, CA2, and CA3, as well as the dentate gyrus (DG). Other methods failed to achieve accurate identification in this region. Finally, for the cortical regions shared by two slices, SpaBatch is able to comprehensively identify and accurately distinguish different subdomains, including the molecular layer (layer 1), external granular and pyramidal layers (layer 2/3), internal granular layer (layer 4), internal pyramidal layer (layer 5), and polymorphic layer (layer 6a and layer 6b), which align with the defined spatial domains 15, 16, 18, 19, 25, and 28 (Figure 4C). We calculated the ARI values of each method on Section 1, with SpaBatch (ARI = 0.44) outperforming the other methods (Figure 4D).

In addition, SpaBatch was able to impute gene expression, as demonstrated through domain marker genes. For example, the expression of the *Neurod6* gene aligned with the CA region identified in domain 11, while *Vxn* and *Lamp5* corresponded to domains 15 and 16, representing the layer 2/3 and layers 6a, 6b, respectively. Furthermore, the combination of *Dner* and *Shf* fully represented the cerebellar region identified in domains 19 and 29 (Figure 4E and Supplementary Fig. S6C, D). This example further highlighted ability of SpaBatch to provide a comprehensive depiction of the sagittal mouse brain while mitigating batch effects, a strength that persisted in Section 2 (Supplementary Fig. S7).

To further explore the adult mouse brain at a microscopic level, we collected 35 coronal slices spanning the anterior-posterior (AP) axis (Ortiz et al., 2020), which include manual annotations of 15 clustering types (Kleshchevnikov et al., 2022). These slices cover the entire brain region from the mouse olfactory bulb to the emergence of the cerebellum (Supplementary Fig. S8). Based on our clustering results and manual annotations, we mapped the locations of these coronal slices onto the Allen Brain Atlas reference and sagittal mouse brain slices obtained through SpaBatch (Figure 5A and Supplementary Fig. S9A). This mapping provides a more comprehensive understanding of the spatial distribution and organization of brain regions in different anatomical planes. By analyzing these slices, we are able to observe the gradual changes in the adult mouse brain from a more microscopic perspective at different anterior-posterior locations. The multi-slice joint analysis task is challenging as it requires methods to account for batch effects across dozens of slices and to distinguish subtle variations in spatial domains within the adult mouse brain.

**Figure 5:**
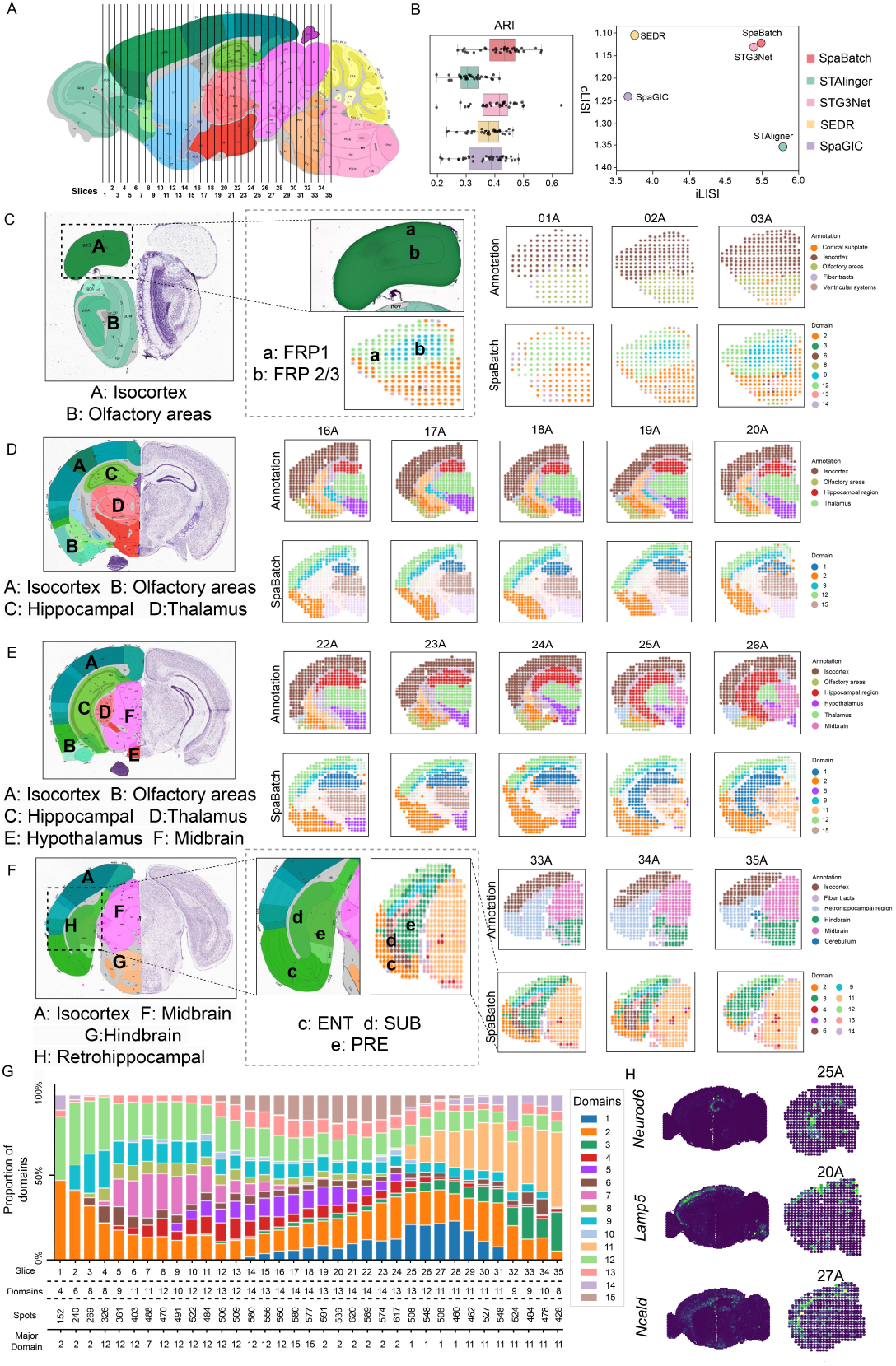
SpaBatch comprehensively depicts the sagittal mouse brain from the micro perspective. A.The locations of the 35 slices were mapped to the Allen Brain Atlas reference based on the spatial domain results identified by SpaBatch and manual annotations. B. The box plot of ARI values computed by SpaBatch and other methods on the 35 coronal mouse brain slices is shown (left). It displays the iLISI and cLISI scores calculated by SpaBatch and other methods on this dataset (right). The x-axis represents the batch mixing score, and the y-axis represents the spatial domain mixing score. The closer the points are to the upper-right corner, the better the performance. C-F. SpaBatch accurately identifies spatial domains in the coronal mouse brain across different slices along the anterior-posterior (AP) axis and corresponds to specific tissue structures through the Allen Brain Atlas reference. SpaBatch can also further identify regions that were not delineated in the manual annotations. G. The stacked bar plot of spatial domain proportions for the 35 slices by SpaBatch. H. The domain marker gene *Neurod6* in the hippocampus and the domain marker genes *Lamp5* and *Ncald* in the cortex exhibit the same spatial pattern in both sagittal and coronal mouse brain slices.

The spatial domain variations identified by SpaBatch across the 35 slices exhibited strong connectivity (Supplementary Fig. S9C, D), and all these variations were validated within the Allen Brain Atlas reference. In the earliest adult mouse brain slices (01A-03A), the regions primarily include the isocortex and olfactory areas. SpaBatch not only accurately identified these two regions but also detected areas that were not differentiated in the manual annotations. Specifically, it precisely subdivided the isocortex into FRP 1 and FRP 2/3 (Figure 5C). As the hippocampus gradually expanded (16A-20A), SpaBatch accurately captured four key spatial domains, which closely aligned with manual annotations (Figure 5D). As the hippocampus continued to expand, the hypothalamus gradually shrank, and the midbrain began to emerge (22A-26A). These dynamic changes were clearly detected by SpaBatch (Figure 5E). As the Hippocampus transitions to Retrohippocampal (33A-35A), SpaBatch divides this area into ENT (entorhinal cortex), SUB (subiculum), and PRE (pre-subiculum) (Figure 5F), which were not differentiated in the manual annotations. These regions play a crucial role in spatial perception, memory formation, and the flow of information between the cortex and the hippocampus (Yao et al., 2021). The clustering proportions of the 35 slices through SpaBatch provided a clear visualization of the spatial domain changes in the mouse brain, such as the emergence of the hippocampus starting from 15A and its disappearance after 31A (Figure 5G). Interestingly, in the sagittal mouse brain slices, SpaBatch captured the same spatial patterns for *Neurod6* expressed in the hippocampus and *Lamp5* and *Ncald* expressed in the cortex within these data sets (Figure 5H). Finally, SpaBatch achieved the best performance in clustering and batch effect correction metrics compared to the baseline methods (Figure 5B and Supplementary Fig. S9B).

In summary, SpaBatch provided a comprehensive depiction of the adult mouse brain’s spatial domains from both macroscopic and microscopic perspectives by analyzing sagittal and coronal mouse brain data. In the sagittal mouse brain, it effectively integrated regions across different brain areas and detected complex structures of cross-slice spatial domains, such as the hippocampus and cerebellar cortex. In the 35 coronal slices spanning the anterior-posterior axis, SpaBatch offered a clear spatial distribution map of mouse brain regions, revealing dynamic changes in structures like the hippocampus and cortex, and linked these to domain-specific marker genes. With its superior spatial domain identification and batch effect correction capabilities, SpaBatch outperformed other methods across multiple evaluation metrics, highlighting its potential as a powerful tool for spatial transcriptomics analysis.

### 3.3 Identifying shared and correlated spatial domains between multiple tissue slices of E9.5 mouse embryos using SpaBatch

To evaluate the performance of SpaBatch across datasets from different platforms, we selected four sections from the E9.5 stage of mouse embryos, profiled using the Stereo-seq platform (Chen et al., 2022), specifically E2S1, E2S2, E2S3, and E2S4. These sections are different slices from the same developmental stage (E9.5), representing the embryonic state at the same time point. During this stage, organogenesis in mice is progressing rapidly, particularly the formation of the heart, nervous system, and major body cavities. Each section covers different regions of the embryo and exhibits anatomical continuity. However, due to spatial positional differences and the presence of batch effects, identifying shared spatial domains across consecutive sections remains a challenge.

Despite the spatial differences in tissue structure and the presence of significant batch effects across these four sections, SpaBatch can effectively integrate the four sections into the embedding space and align the spatial domains (Figure 6A). We compared the performance of SpaBatch with STAligner, STG3Net, SEDR, and SpaGIC in integrating the four consecutive tissue sections. Our analysis indicated that SpaBatch was the only method capable of fully identifying the heart region in all four sections. It demonstrated exceptional ability to align spatial domains, effectively integrating data from all four sections. In contrast, STAligner emerged as the most competitive method, showing excellent performance in aligning and integrating tissue sections, although it could not fully capture the heart region across all slices as SpaBatch did. SpaBatch not only comprehensively identified the heart region in the mouse embryo but also excelled in aligning other tissues. It successfully recognized additional spatial domains across the four consecutive sections, including Mesenchyme (domain 8), Sclerotome (domain 9), Primitive gut tube (domain 10), Brain (domain 11) and Spinal cord (domain 12) (Supplementary Fig. S10A-C). SpaBatch shows significantly better ARI, ACC, and V-measure on the four slices compared to the baseline methods, indicating its more precise recognition of spatial structures (Figure 6D). This ability to accurately capture and align various tissue structures across different spatial domains further highlights the robustness and versatility of SpaBatch in handling complex and heterogeneous spatial transcriptomics data, enabling a comprehensive understanding of the tissue organization within the developing embryo. We further explored the relationships between different spatial domains using a spatial domain correlation heatmap and observed a significant correlation between domain 1 and domain 13 (Figure 6B, C). This finding drew our attention, as the close connection between these two domains may reveal their potential functional association during mouse embryonic development. In subsequent analysis, we compared these two domains with manually annotated regions in the embryo and found that domain 1 and domain 13 strongly overlapped with the annotated heart and cavity regions. The marker gene *Sh3bgr* exhibited significant and specific expression in this region across all fourslices, further supporting their correlation (Figure 6E). *Sh3bgr* has been reported to be associated with myocardial development (Deshpande et al., 2021). Moreover, the associated regions also include domain 11 and domain 12. From the correlation heatmap, it was observed that these two spatial domains highly overlap with the manually annotated brain and spinal cord (Supplementary Fig. S10A). We performed GO enrichment analysis on the two spatial domains. The spatial domain highly associated with the brain primarily includes neuron development, synapse organization, axon guidance, and other functions, which are closely related to the complex neural network development and functional regulation in brain tissues. In the spatial domain highly associated with the spinal cord, similar neuron function-related terms also appeared, such as neuron migration, axon repair, and motor neuron regulation. There is significant functional overlap in the GO enrichment results of these two spatial domains, demonstrating highly consistent functional directions (Supplementary Fig. S10D). This overlap and similarity in terms validate the high correlation between domain 11 and domain 12 in the correlation heatmap.

**Figure 6:**
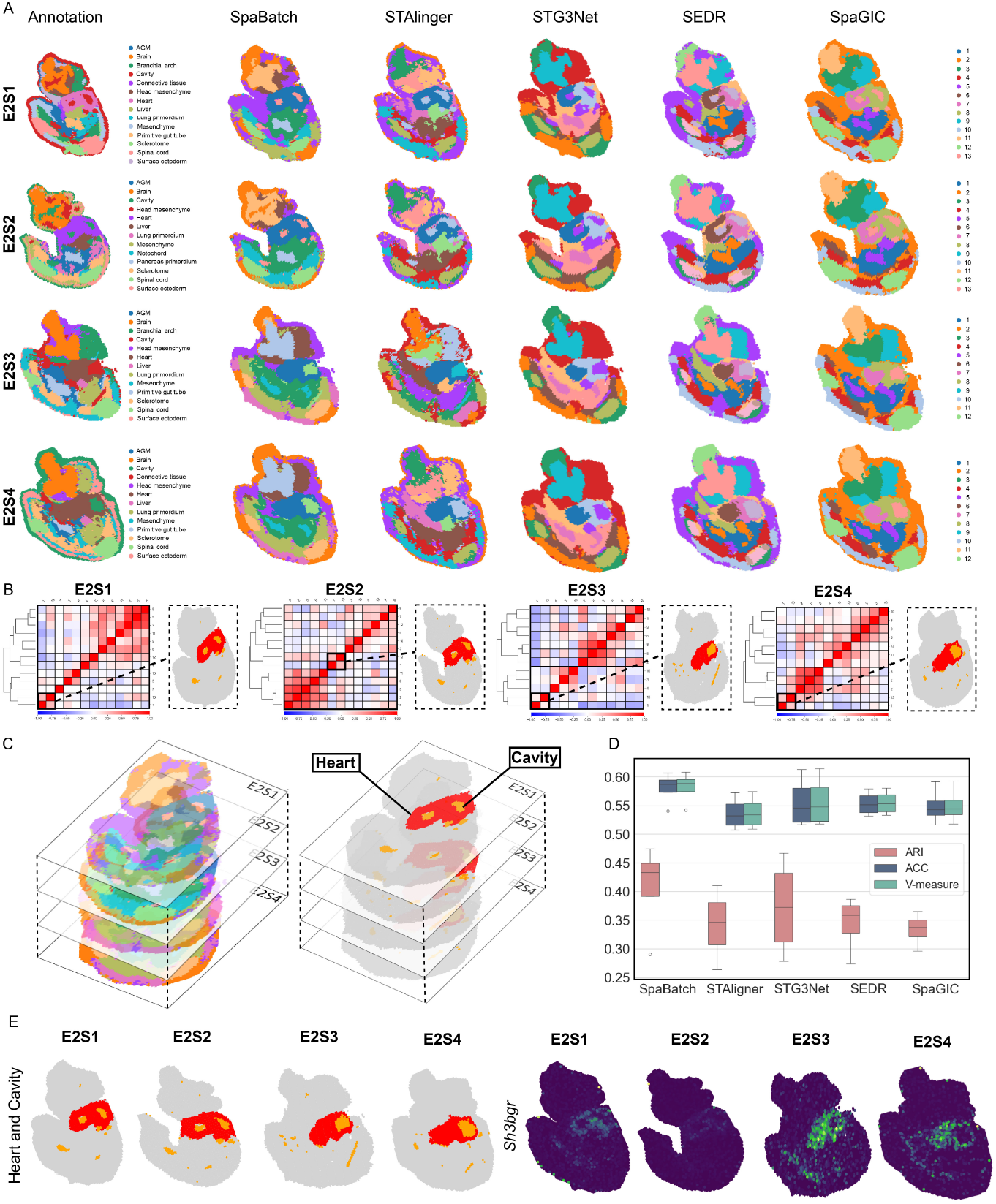
SpaBatch cross-slice matching of shared spatial domains on the mouse embryo dataset. A. Spatial domains identified by SpaBatch and other methods on the E2S1, E2S2, E2S3, and E2S4 slices of the E9.5 embryo. B. SpaBatch identifies correlated spatial domains 1 and 13 across the E2S1, E2S2, E2S3, and E2S4 slices of the E9.5 embryo through a correlation heatmap, and these domains highly overlap with the manually annotated heart and cavity regions. C. Spatial domains were identified by SpaBatch across all four slices, E2S1, E2S2, E2S3, and E2S4 (left). The regions of correlation between the heart and cavity identified by SpaBatch on all four sections (right). D. Box plots of ARI, ACC, and V-measure calculated by SpaBatch and other methods on the mouse embryo E2S1, E2S2, E2S3, and E2S4 slices. E. SpaBatch identified spatial domains related to the heart and cavity across all four slices (left), along with the spatial expression of the marker gene *Sh3bgr* associated with this region (right).

### 3.4 SpaBatch identifies the development of the human heart

We applied SpaBatch to the human heart ST dataset, which includes two sets of slices collected from human embryonic hearts at 4.5-5 and 6.5 PCW (Figure 7A). This data was used to explore SpaBatch’s ability to identify the dynamic changes in tissue structures across slices with developmental processes.

**Figure 7:**
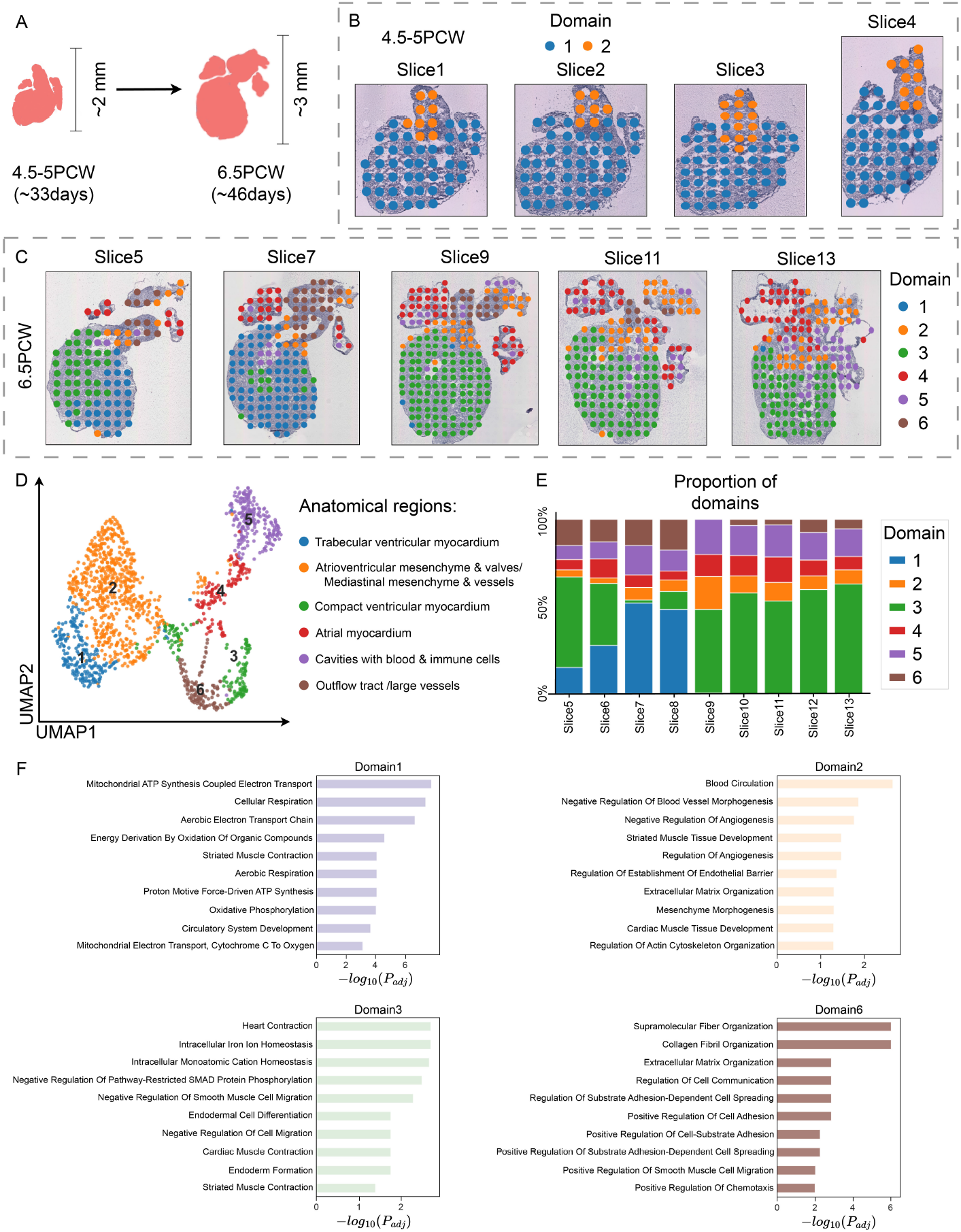
SpaBatch identifies the development of the human heart. A. The development of the human heart from 4.5-5 PCW to 6.5 PCW. B. SpaBatch spatial domain identification in the human heart at 4.5-5 PCW. C. SpaBatch spatial domain identification in the human heart at 6.5 PCW. D. UMAP visualization of the human heart at 6.5 PCW and its corresponding anatomical regions. E. SpaBatch spatial domain proportion stacked bar plot in the human heart at 6.5 PCW. F. GO enrichment analysis of spatial domains in the 6.5 PCW heart.

At 4.5-5 PCW, SpaBatch identified two distinct spatial domains, which further developed at 6.5 PCW (Figure 7B). Next, we focused on the human heart at 6.5 PCW, where SpaBatch identified six spatial domains across nine sections Figure 7C and Supplementary Fig. S11). Based on the anatomical region annotations provided by a previous report (Asp et al., 2019), we mapped the identified spatial domains to their corresponding anatomical regions (Figure 7D). For example, spatial domains 1, 3, and 4 corresponded to the trabecular ventricular myocardium, compact ventricular myocardium, and atrial myocardium, respectively. These were all important components of the cardiac muscle. The trabecular myocardium and compact myocardium were typically present in the ventricles. The compact myocardium provided strong contractile force, while the trabecular myocardium was responsible for structural support and local blood flow. The atrial myocardium supported the overall blood flow of the heart and coordinated the contraction between the atria and ventricles (Meilhac and Buckingham, 2018). By visualizing the proportion of spatial domains identified by SpaBatch, the dynamic changes in the 6.5 PCW human heart tissue could be intuitively observed (Figure 7E).

The human heart data did not have manual annotations as ground truth. For the identified spatial domains, we performed GO enrichment analysis to characterize their functions (Fang et al., 2023). Domain 1 corresponds to the trabecular ventricular myocardium region, and the top enriched GO terms are primarily related to cellular respiration. The GO terms enriched in domain 2 involve blood circulation, vasculature morphogenesis, angiogenesis, etc., which align with its corresponding anatomical region. Domain 3 corresponds to the compact ventricular myocardium region and is enriched with processes related to cardiac development and heart contraction. The terms enriched in domain 6 are associated with extracellular matrix (ECM) tissue and are related to the integrity and stability of the cardiovascular system (Figure 7F).

### 3.5 Leveraging limited annotations for accurate spatial domain detection in the HER2-positive breast cancer dataset

In addition to normal tissue, we applied SpaBatch to the HER2-positive breast cancer dataset to evaluate its ability to make biological discoveries in abnormal tissue slices. The dataset consists of tumor samples from 8 different patients, represented as groups A-H. Groups A-D each contain 6 slices, while groups E-H each contain 3 slices. We performed separate analyses for each of the A-H groups (Andersson et al., 2021; Wu et al., 2021).

In groups A-H, only the first slice was annotated by a pathologist. We conducted experiments with SpaBatch, STAligner, STG3Net, SEDR, and SpaGIC on this dataset, and calculated the ARI on the first slice (Supplementary Fig. S12). As shown in Figure 8A, SpaBatch outperformed other methods in spatial clustering performance across all slices in all groups. In terms of overall clustering performance, SpaBatch also achieved the highest median (ARI = 0.367) and mean (ARI = 0.362), outperforming the second-place STAligner with a median (ARI = 0.196) and mean (ARI = 0.255), showing a significant improvement (Figure 8B). We then focused on the H group data, which contains more diverse spatial domains. Neither SpaGIC nor SEDR were able to identify the cancer in situ region. STG3Net failed to correctly identify the breast glands. SpaBatch and STAligner performed the best on slice H1, correctly identifying continuous regions such as cancer in situ and adipose tissue, as well as discrete regions such as breast glands. SpaBatch outperformed STAligner in spatial domain range and boundary identification, achieving a higher ARI (Figure 8C). Interestingly, SpaBatch adjusted the training on labeled data and predicted the unlabeled data within the same group through joint analysis. By using the manual annotation on the first slice and the H&E images of the remaining slices, excellent results were observed in slices H2 and H3. In the spatial domainidentification results for H2 and H3 slices using SpaBatch, domain 6 and the breast glands region aligned very well, and the detected cancer regions (cancer in situ and invasive cancer) matched the dark areas in the H&E images. This approach effectively utilized limited annotated data for a semi-supervised learning-like process, enabling efficient spatial transcriptomics analysis without fully annotated data (Figure 8D).

**Figure 8:**
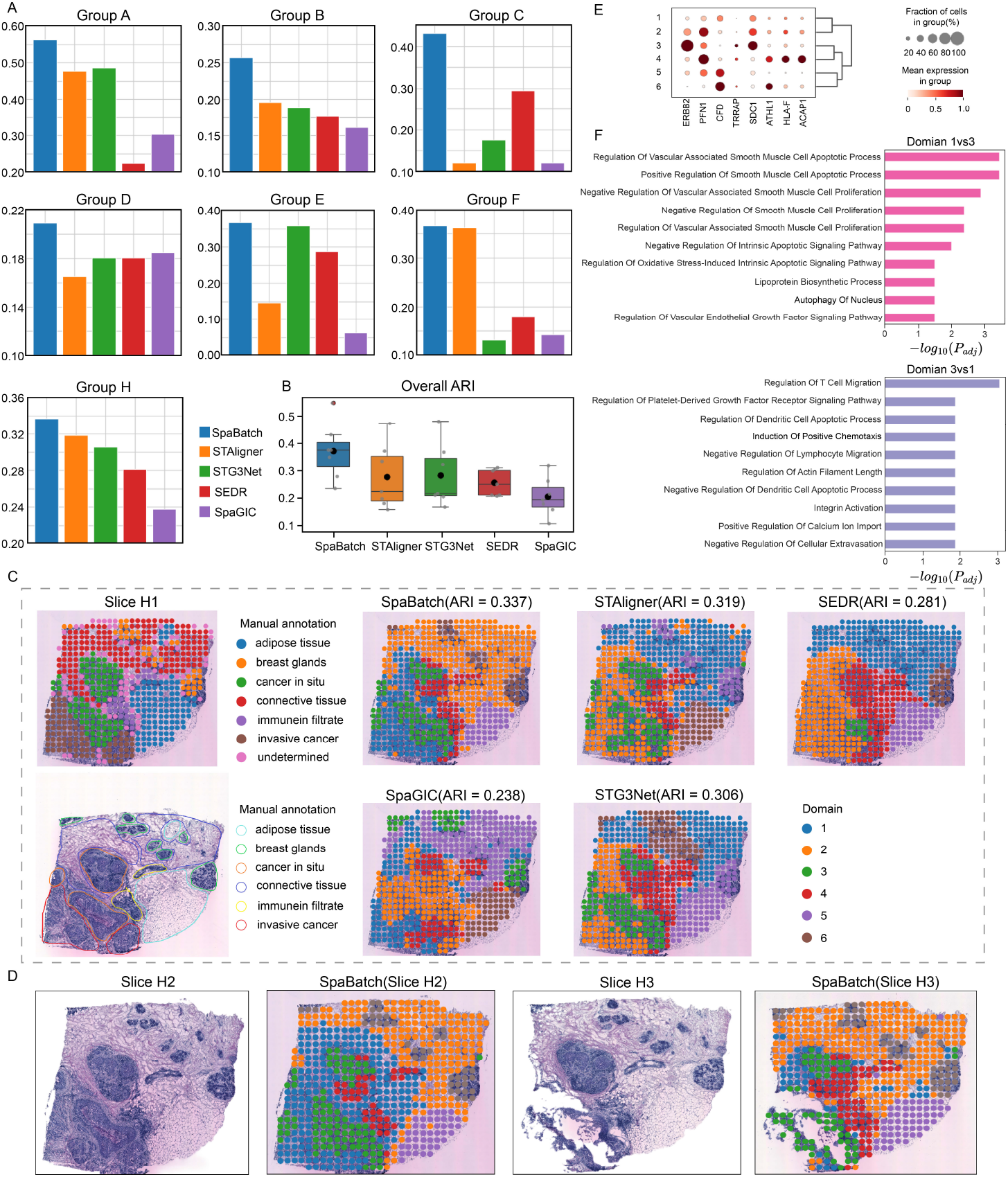
Leveraging limited annotations for accurate spatial domain detection in the HER2-positive breast cancer dataset. A. The ARI calculated for the first slice of group A-H in the HER2-positive breast cancer dataset using SpaBatch compared with other methods. B. The overall boxplot of ARI values for each group. C. Manual annotation of slice H1 (left), and the results of spatial domain identification on slice H1 using SpaBatch and other methods (right). D. The spatial domains of slices H2 and H3 predicted by SpaBatch through joint training on the H group data. E. Bubble plot of the differential analysis between cancer regions and other regions. F. GO enrichment analysis results of the in situ cancer and invasive cancer.

We performed differential analysis between cancer regions and other regions. As shown in Figure 8E, SpaBatch indeed identified that the cancer in situ region recovered from domain 3 highly expresses the breast cancer marker *ERBB2* compared to other regions (Fernandez et al., 2022). Next, we investigated the heterogeneity of the cancer regions. Spatial domain 3 exhibited various immune response-related GO terms, including signaling pathways for B cells and T cells, as well as the regulation of respiratory burst, suggesting a strong immune response in the in situ cancer region. On the other hand, spatial domain 1 was enriched in pathways that inhibit cell apoptosis, which may indicate the presence of more anti-apoptotic mechanisms in invasive cancer, helping tumor cells evade apoptosis (Jan et al., 2019). The differences in enriched biological processes highlight the distinctions between these two regions, emphasizing their respective immune/stromal microenvironments (Figure 8F).

## 4 CONCLUSION

In this paper, we present a deep learning framework, SpaBatch, to remove batch effects from multi-slice spatial transcriptomics data. By leveraging Variational Graph Autoencoders, self-supervised learning, and triplet learning with readout aggregation, SpaBatch enables robust cross-slice alignment and enhances the analysis of spatial transcriptome domains. Validated across diverse tissue types and experimental conditions, SpaBatch demonstrates its potential to uncover new biological insights and improve clustering accuracy in multi-slice spatial transcriptomics studies. This framework paves the way for more comprehensive and integrative analyses in spatial biology.

## Acknowledgements

The work was supported in part by the National Natural Science Foundation of China (62262069).

